# Embracing scale-dependence to achieve a deeper understanding of biodiversity and its change across communities

**DOI:** 10.1101/275701

**Authors:** Jonathan M. Chase, Brian J. McGill, Daniel J. McGlinn, Felix May, Shane A. Blowes, Xiao Xiao, Tiffany M. Knight, Oliver Purschke, Nicholas J. Gotelli

## Abstract

Because biodiversity is multidimensional and scale-dependent, it is challenging to estimate its change. However, it is unclear (1) how much scale-dependence matters for empirical studies, and (2) if it does matter, how exactly we should quantify biodiversity change. To address the first question, we analyzed studies with comparisons among multiple assemblages, and found that rarefaction curves frequently crossed, implying reversals in the ranking of species richness across spatial scales. Moreover, the most frequently measured aspect of diversity—species richness—was poorly correlated with other measures of diversity. Second, we collated studies that included spatial scale in their estimates of biodiversity change in response to ecological drivers and found frequent and strong scale-dependence, including nearly 10% of studies which showed that biodiversity changes switched directions across scales. Having established the complexity of empirical biodiversity comparisons, we describe a synthesis of methods based on rarefaction curves that allow more explicit analyses of spatial and sampling effects on biodiversity comparisons. We use a case study of nutrient additions in experimental ponds to illustrate how this multi-dimensional and multi-scale perspective informs the responses of biodiversity to ecological drivers.

**Statement of Authorship:** JC and BM conceived the study and the overall approach, and all authors participated in multiple working group meetings to develop and refine the approach. BM collected the data for the meta-analysis that led to Fig. 2,3; JC collected the data for the metaanalysis that led to Figure 4 and S1; SB and FM did the analyses for Figures 2-4; DM, FM and XX wrote the code for the analysis used for the recipe and case study in Figure 6. JC, BM and NG wrote first drafts of most sections, and all authors contributed substantially to revisions.

Figure 1.
A. Individual-based rarefaction curves of three hypothetical communities (labelled A,B, C) where ranked differences between communities are consistent across scales. B. Individual-based rarefaction curves of three hypothetical communities (labelled A,B, C) where rankings between communities switch because of differences in the total numbers of species, and their relative abundances. Dotted vertical lines illustrate sampling scales where rankings switch. These curves were generated using the sim_sad function from the mobsim R package (May et al. 2018).

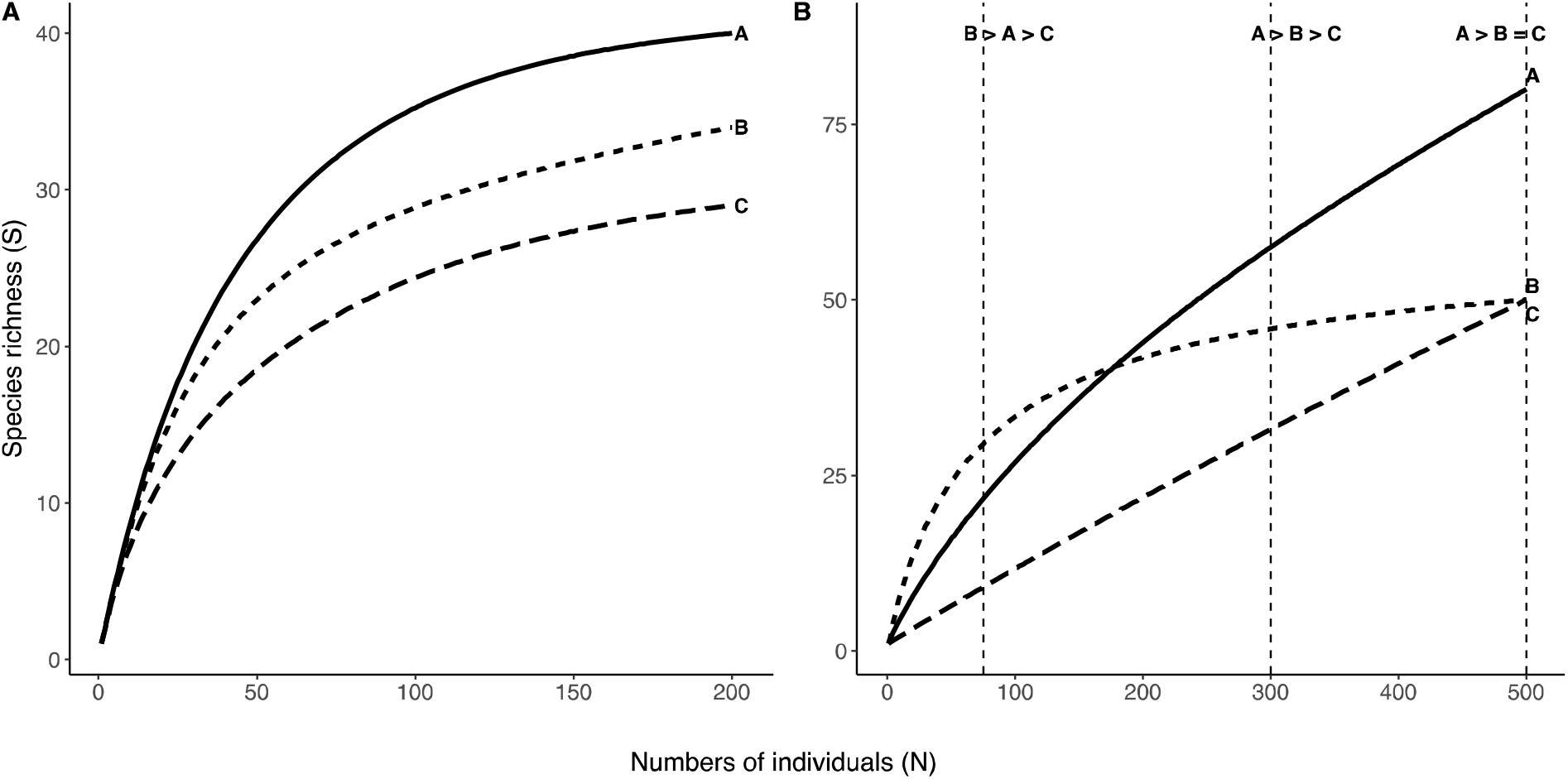

Figure 2.
Bivariate relationships between N, S_PIE_ and S for 346 communities across the 37 datasets taken from McGill (2011b)(see Appendix 1). (A) S as a function of N; (B) S as a function of S_PIE_. (N vs S_PIE_ not shown). Black lines depict the relationships across studies (and correspond to R^2^ fixed); colored points and lines show the relationships within studies. All axes are log-scale. Insets are histograms of the study-level slopes, with the solid line representing the slope across all studies. Gray bars indicate the study-level slope did not differ from zero, blue indicates a significant positive slope, and red indicates a significant negative slope.

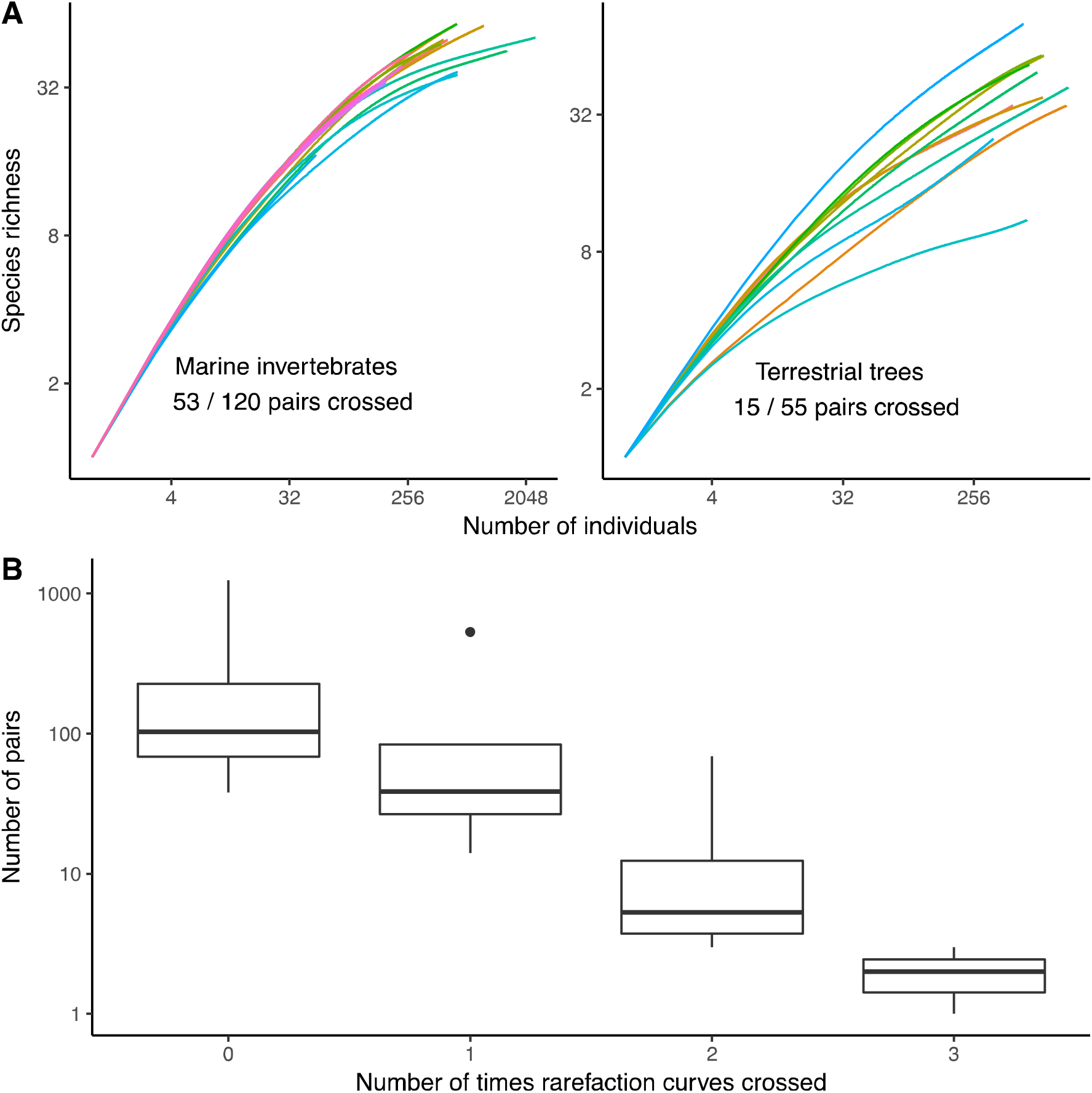

Figure 3.
Representative rarefaction curves, the proportion of curves that crossed, and counts of how often curves crossed. (A) Rarefaction curves for different local communities within two datasets: marine invertebrates (nematodes) along a gradient from a waste plant outlet (Lambshead 1986), and trees in a Ugandan rainforest (Eggeling 1947); axes are log-transformed. (B) Counts of how many times pairs of rarefaction curves (from the same community) crossed; y-axis is on a log-scale.

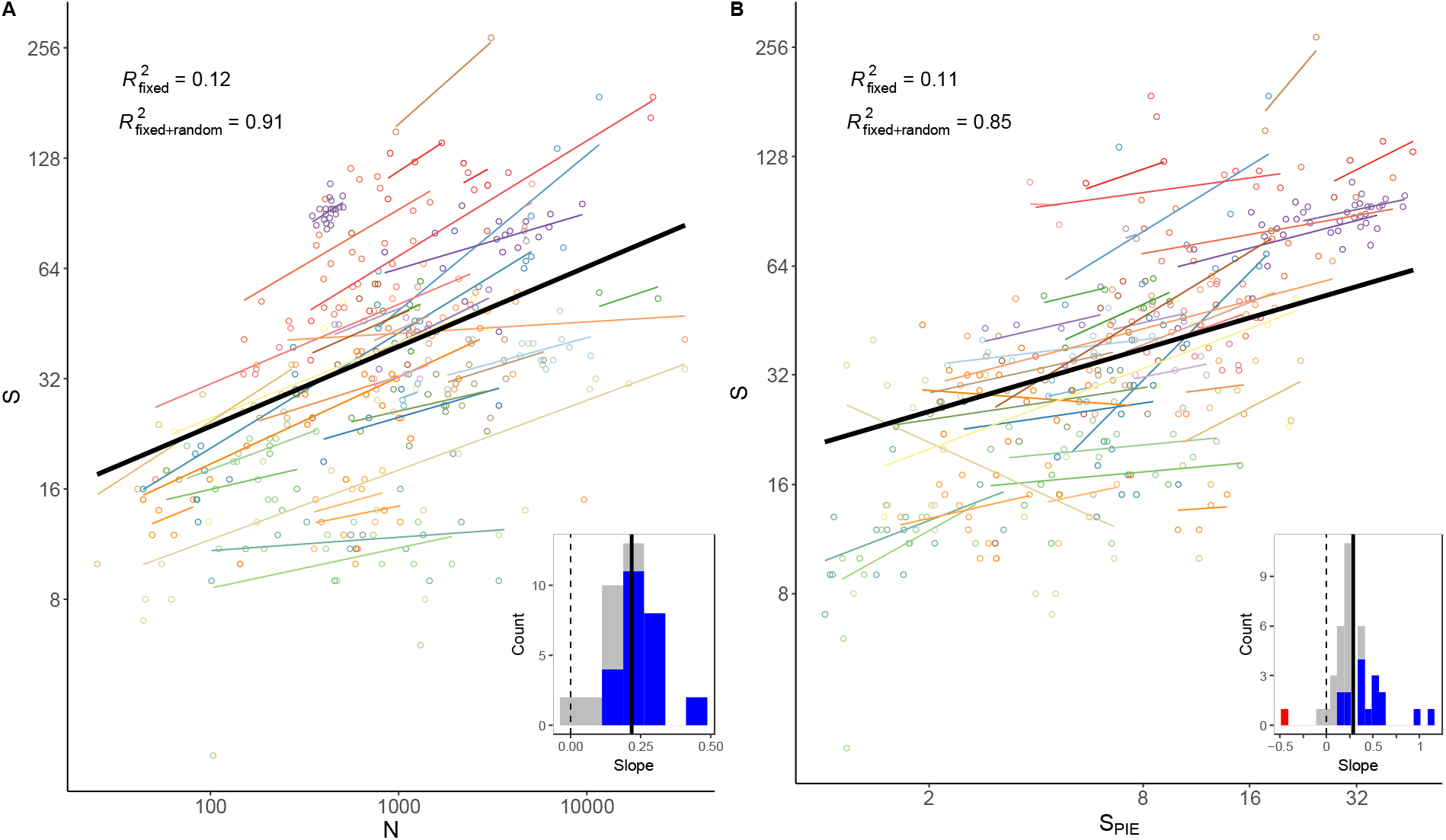

**Data accessibility statement:** All data for meta-analyses and case study will be deposited in a publically available repository with DOI upon acceptance (available in link for submission).

## Introduction

Biodiversity is multidimensional and scale-dependent. It is multi-dimensional, because it is determined by a combination of the total numbers of species in a given area and the total and relative abundances of those species in that area. And it is scale-dependent, because all measures of diversity necessarily increase non-linearly with increasing sampling effort and spatial scale. However, the nature of this scaling depends on the total numbers of species, the total and relative abundances of individuals, as well as their spatial distribution. This makes comparisons of biodiversity from place to place (and time to time) more challenging than comparisons of most other variables in ecology (e.g., McGill 2011a).

Understanding how biodiversity changes in the face of natural and anthropogenic drivers can be greatly enhanced by taking a scale-explicit and multi-dimensional perspective on biodiversity and its change. Of course, ecologists have long recognized the complexity in making comparisons of biodiversity in response to natural and anthropogenic drivers and continuously sought better solutions to this problem (e.g., Preston 1960, MacArthur and MacArthur 1961, Hurlbert 1971, Hill 1973, Lande 1996, Gotelli and Colwell 2001, Jost 2006, Magurran and McGill 2011, Chao and Jost 2012, Chase and Knight 2013). Nevertheless, there is little consensus or consistency as to which (if any) solutions are best. And as a likely result, the vast majority of empirical studies ignore these complexities and typically measure and compare only a single variable (usually species richness), measured at a single spatial scale (usually determined by the grain size and extent of the sampling).

To illuminate the neglect of spatial scale and multi-dimensionality in biodiversity studies, we evaluated papers published from 2012-2017 in Ecology Letters that compared (taxonomic) diversity (but not compositional). Of these, 80% (44/55) only reported one measure of diversity (90% of those were species richness), and 75% (41/55) compared diversity at only one spatial scale; just three exemplary studies (Solar et al. 2015, Martinson et al. 2017, Livne-Luzon et al. 2017), measured multiple dimensions of diversity at multiple scales, which we advocate to more fully understand biodiversity change.

Simply estimating a difference (i.e., effect size) of any measure of diversity at a single scale and then subjecting that difference to standard statistical analyses does not capture the fact that the magnitude, and even direction, of biodiversity change critically depends on which measure, and at which spatial scale, the data were collected (e.g., Whittaker et al. 2001, Chase and Leibold 2002, Rahbek 2005, Dumbrell et al. 2008, Sandel and Smith 2009, Keil et al. 2011). This is not just a theoretical issue, but is a concrete problem that can affect biodiversity conservation and management. For example, Hill and Hamer (2004) reviewed studies of the effects of logging on tropical forest bird diversity and found that bird diversity tended to decrease with logging when the scale in which diversity was measured was relatively small (i.e., less than 25 ha), but increase with logging in studies in which diversity was measured at larger spatial scales. In an analysis and review of the effects of logging on butterfly diversity, Hamer and Hill (2000) likewise illustrated scale-dependent effects of logging, but in a different direction (i.e., minor effects of logging on diversity at small scales, but larger negative effects at larger scales).

Does it matter that biodiversity is multidimensional, where change in an estimate of species richness at a given scale might differentially reflect changes in the total and relative abundances of individuals in a community? Many have argued that different measures of biodiversity (e.g., evenness, richness) are highly correlated and thus perhaps redundant. If so, one measure of diversity and its change would likely suffice. Likewise, does it matter that these biodiversity changes can be scale-dependent? While there are certainly many case studies where scale-dependence has been observed, is this common? How often do measured effect sizes vary in magnitude and/or direction such that conclusions about biodiversity change would be altered?

Here, we first review the multidimensional concept of biodiversity and its scaling to establish general hypotheses about the scale-dependent nature of biodiversity measures and their comparisons. We collated datasets where species assemblages from multiple sites were compared within a study and asked whether measures of biodiversity were correlated. Next, we collated studies in which biodiversity changes were measured at more than one spatial scale, so that we could estimate how frequent scale-dependence occurred in the magnitude and direction of effect sizes. Finally, after showing that biodiversity changes are indeed typically multivariate and scale-dependent, we describe a recipe that synthesizes a number of measures of biodiversity to more fully evaluate how biodiversity changes, and we illustrate our approach with a case study from a previously published experiment on aquatic mesocosms.

## The multidimensional nature of biodiversity and its scaling

Biodiversity is a summary variable that is influenced by the abundances and distributions of populations of multiple species. Three factors in particular combine to influence diversity measures at any given sample or scale (e.g., He and Legendre 2002, McGlinn and Palmer 2009, McGill 2011a, Chase and Knight 2013): (1) the total numbers of individuals (N) in a community; (2) the Species Abundance Distribution (SAD), which includes both the total numbers of species in the community (S) and their relative abundances; (3) the spatial distributions of individuals, which influences patterns of intraspecific aggregation or clumping. Each of these components can be independently and interactively influenced by ecological drivers, leading to multidimensional and scale-dependent biodiversity change. We recognize that a critical component of biodiversity change is in compositional shifts, which uses a different set of metrics to detect changes (e.g., Baselga 2010, Dornelas et al 2014, Legendre 2014, Hillebrand et al. 2018). But here we only focus on the quantifications of biodiversity within and across communities, which presents more than enough challenge in itself.

When comparing communities, any differences in the N, the SAD and/or aggregation will influence how measurements and comparisons of biodiversity change with sampling scale or effort. Figure 1 illustrates one way to depict this scaling—via individual-based rarefaction curves— and shows two qualitatively distinct ways in which it can differ. Differences in individual-based rarefaction curves are determined only by differences in the SAD (both the total S and their relative abundance) (e.g., Gotelli and Colwell 2001, Cayuela et al. 2015)

In Figure 1A, three communities (labelled A, B, and C) differ in a straightforward way; both species richness and the relative abundances of the three communities are ranked A>B>C. Thus, if we were to compare species richness or other commonly measured estimates of diversity (e.g., Shannon, Simpson), they would rank the same. In such a case, it would not be necessary to worry much about the multidimensionality of diversity estimates (i.e., because different dimensions are correlated). However, even when the rankings of S and other diversity measures are consistent, there can still be scale-dependence when the SAD changes among communities (e.g., Gotelli and Colwell 2001, Cao et al. 2007, Chase and Knight 2013, Cayuela et al. 2015). Nevertheless, this scale-dependence is moderate and would be predictable, consistently leading to larger differences between communities as scale increases (i.e., rarefaction curves diverge).

In Figure 1B, three communities (labelled A, B, and C) differ in a more complex way. Community A is dominated by a few very common species and has several rarer species (i.e., is less even), and thus has a shallower slope near its base than Community B, but higher total S. That is, the difference between the communities flip-flop; at small scales, B is richer than A, but at large scales, A is richer than B. The comparison between Communities B and C is somewhat similar; they have the same total S, but different levels of evenness (and diversity), again decoupling the different measures of diversity. Finally, Communities A and C differ in both S and diversity measures in the same direction.

These represent two extreme scenarios for diversity comparisons between communities. In Figure 1A, diversity measures are largely correlated and scale-dependence is unidirectional, suggesting that we may not need to worry much about which measures of diversity we use for comparisons, and at which scales. Alternatively, in Figure 1B, diversity measures are decoupled and scale dependence is strong, suggesting that more complex measurements and comparisons of biodiversity will be necessary. Furthermore, the curves in Figure 1 only include differences in the SAD between communities, whereas variation also in N and aggregation will lead to even more complex SACs, which could further decouple diversity measures and create more, and more complex, forms of scale-dependence (see also Chase and Knight 2013, Blowes et al. 2017, McGlinn et al. 2018).

Next, we empirically evaluate these possibilities, with the extremes being: (i) biodiversity change between communities is relatively “well behaved”, where different measures of biodiversity are largely correlated and scale-dependence is moderate and unidirectional (Figure 1A), or (ii) biodiversity change is more complicated where different measures are poorly correlated and scale-dependence is strong and multi-directional (Figure 1B). As discussed above, few exemplary data exist where multiple measures and multiple scales of biodiversity change are collected and compared. And so we used two separate meta-analyses described in the following sections, one to compare correlations (or lack thereof) among diversity measures, and another containing data measured at multiple scales to evaluate the degree and direction of scale-dependence in biodiversity responses to ecological drivers.

## Empirical evidence that biodiversity change is often multidimensional

To determine which of the caricatures in Figure 1 is more indicative of biodiversity change among communities, we directly compared SADs and the resulting rarefaction curves from 37 datasets in which two or more local communities were compared (data and references listed in Appendix 2, taken from a larger assemblage of datasets amassed by McGill 2011b). We used these studies to evaluate (i) how often rarefaction curves cross, suggesting that sampling scale can critically influence conclusions about the magnitude and direction of biodiversity change, and (ii) how measures of diversity (S, S_PIE_) and numbers of individuals (N) co-vary among communities. Because there was no spatial information available in these datasets, we were only able to compare rarefaction curves, eliminating potential influences of differences in spatial aggregation, which could have further complicated their relationships.

Example datasets include bird surveys along standardized routes in North America (i.e., the Breeding Bird Survey; Pardieck et al. 2017), benthic marine nematodes collected from cores in relation to pollution (Lambshead 1986), arthropods associated with different macrophyte species in streams (Harrod 1964), and trees from different sections of the Barro Colorado Island permanent forest plot (Condit et al. 2012). Most datasets compared 2-20 communities; for datasets with >20 communities (e.g., the North American breeding birds), we randomly selected 20 sites so that larger datasets did not numerically dominate the results.

First, we evaluated whether the rankings of rarefaction curves were relatively well-behaved, as in Figure 1A, or whether curves were more likely to cross, as in Figure 1B (see also Lande et al. 2000, Thompson and Withers 2003). Such crossings indicate not just a change in magnitude, but also a change in direction of biodiversity change due to sampling and spatial scale. We compared curves for each pair of communities within a single dataset (see Figure 2A for two examples). Across all datasets, a total of 2203 pairwise comparisons were made, with 732 pairs of rarefaction curves crossing (33% of all pairs, with some curves crossing multiple times) (Figure 2B). Importantly, the crossing points were not ecologically trivial (e.g., crossings only at very small N). The average crossing occurred at 24% of the total abundance in a community and the average vertical separation on curves that crossed was 6.3 species (versus an average of ~40 species in a community).

Next, we examined the relationships between three measures often collected and compared among communities. Specifically, we compared the total N measured in a community, the number of species (S) from that community, and a measure of diversity that takes relative abundances into account. For the latter, we used a bias-corrected version of Hurlbert’s (1971) Probability of Interspecific Encounter; 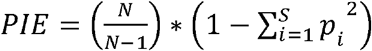; where N is the total number of individuals in the entire community, S is the total number of species in the community, and p_i_ is the proportion of each species i. We used PIE because it is a relatively unbiased estimator of evenness, it is intuitive and familiar to ecologists, and is equivalent to the slope of the rarefaction curve at its base (between N=1 and N=2) (Olszewski 2004). Specifically, a more even community (higher PIE) accumulates species more quickly with increasing N than a less even community (lower PIE). PIE is equivalent to 1-Simpson’s index (also called the Gini-Simpson index) (Jost 2006), but we prefer the PIE terminology because it has a strong intuitive interpretation (i.e., the probability of interspecific encounter), as well as connection to the individual-based rarefaction curve. We converted PIE into an effective number of species, S_PIE_, so that comparisons with S are more intuitive (e.g., Hill 1973, Jost 2006, 2007); S_PIE_ is equivalent to 1/Simpson’s index. S_PIE_ is the number of equally abundant species it would take to yield a given value of PIE (see McGlinn et al, 2018 for justification for why we take the effective number of species from the bias corrected PIE). In a community in which all species had identical abundances, S_PIE_ would be equal to S, but as the community becomes increasingly dominated, S stays the same, but PIE decreases towards 0 and S_PIE_ decreases towards 1. Other measures of diversity when converted to an effective number of species are also potentially informative, such as Shannon’s index (e.g., Jost 2006, 2007, Chao et al. 2014a), but here we focus on S and S_PIE_ because S is most sensitive to changes in the abundance of rare species, whereas PIE is more sensitive to changes in the abundance of common species.

We examined bivariate relationships between S, N and S_PIE_ using hierarchical linear models with a fixed effect, and modeled variation among studies as a random effect on slopes and intercepts. N and S were modelled assuming Poisson error and a log-link function, and S_PIE_ was log-transformed and modelled assuming Gaussian error; N, S_PIE_ and S entered all models as log-transformed covariates. All models were fit in R (R Development Core Team 2017) using the brms package (Bürkner 2017). To compare variation across versus between datasets, we quantified the marginal (fixed effect only) and conditional *R*^2^ (fixed + random effects; Gelman et al. 2017).

First, we regressed S as a function of N (Figure 3A). As expected from sampling, there was an overall increasing relationship between N and S across all studies. However, the relationship had low explanatory power (*R*^2^ = 0.12). The variation explained when the relationships were allowed to vary between studies increased dramatically (*R*^2^ increases to 0.91). While most studies showed a positive relationship between N and S, slopes and intercepts among studies varied widely, and more than 30% (12/37) of studies showed that the 95% credible interval of the study-level slope overlapped zero. Clearly, much of the relationship between N and S, as well as its variation among studies, is due to sampling and methodological differences. However, the fact that some types of communities within a study have large differences in N, with little variation in S, whereas other communities within a study have smaller N differences, but large S differences indicates that N and S changes are not completely coupled. For example, there are many instances where N can dramatically increase or decrease from one place to another, but if there is a concomitant change in the relative abundances of species, we might expect little variation in S.

Second, we regressed S as a function of S_PIE_ to determine whether diversity measures were consistent in their differences among communities (Figure 3B). Again, we found a significant, but weak (*R*^2^ = 0.11), relationship between these two measures. As in our analysis of S and N, the fit of the model was much higher when we incorporated a random effect to capture variation in the slope and intercept among studies (*R*^2^ = 0.85). Here, most studies (20/37) showed that the 95% credible interval of the study-level slope overlapped zero, though some were positively associated (16/37) and one showed negative associations.

In sum, our analyses above show that rarefaction curves often cross and that different measures among communities are often decoupled (N, S, S_PIE_). This supports the hypothesis that multiple measures are often necessary to understand treatment effects on biodiversity (see also e.g., Stirling and Wilsey 2001, Wilsey et al. 2005, Soininen et al. 2012).

## Empirical evidence that biodiversity change is often scale-dependent

Explicit consideration of spatial scale has provided some resolution to contradictory results in the literature. For example, Chase and Leibold (2002) showed that the influence of productivity on species richness was much stronger at larger spatial scales than at smaller ones (see also e.g., Whittaker et al. 2001, Gardezi and Gonzalez 2008, Andrew et al. 2012). Likewise, Powell et al. (2011, 2013) showed that invasive species can simultaneously have strong impacts on diversity at small scales, but much weaker effects at larger scales.

Although such multiple-scale analyses remain uncommon in the biodiversity literature, there are enough to allow for a synthetic meta-analysis. We identified possible studies using an ISI Web of Science search with the key words “species richness or diversity or biodiversity” AND “scale or grain or extent” AND “ecology” (to eliminate hits that were in other fields). This yielded ~8,500 papers, from which we could extract species richness at more than one scale in response to ecological treatment from 103 comparisons within 52 studies (several studies reported responses from more than one driver, taxonomic group, or study site). A list of the studies and their data are given in the Supplemental material (Appendix 2).

With this collection of studies, we asked how frequent, and in which direction, the change in biodiversity was influenced by spatial scale. We coded the log ratio effect size for any case where species richness was lower in the treatment than the control as negative, and any case where species richness was higher in the treatment than in the control as positive. Unfortunately, we do not have estimates of uncertainty for most studies (i.e., most studies did not provide an estimate of variance at the largest scale, and it was not possible to extract variance for many smaller scale studies), and so cannot explore potential publication bias using traditional approaches (e.g., funnel plots). We further note that there is likely some bias because our search criteria explicitly included ‘scale’ (or ‘grain’, ‘extent’) as a search term. Thus, our meta-analysis should be taken as a first indication of the frequency and direction of scale-dependent biodiversity change.

The analysis shown in Figure 4 shows that scale-dependence in frequent, and a non-trivial modifier of effect sizes in empirical studies. Most studies showed consistent positive (upper right quadrant) or negative (lower left quadrant) effects of the treatment at both small and large spatial scales. However, this obscures the reality that there are many instances where strong effects at one scale were weak at the other. If effect sizes were scale-free then all points would fall on the 1:1 line, but this line explains only 30% of the variance. Furthermore, nearly 12% of comparisons (12/103) showed a reversal in direction of species richness change from negative to positive (upper left quadrant), or from positive to negative (lower right quadrant) from smaller to larger scales, several of which are quite substantial. While this is lower than the 33% of crossing rarefaction curves (implying reversals) that we found in the analysis above (Figure 3), it again implies that biodiversity change can not only vary in magnitude, but can often switch direction. Overall, effect size at one scale is a poor predictor of effect size at a different scale.

**Figure 4.**
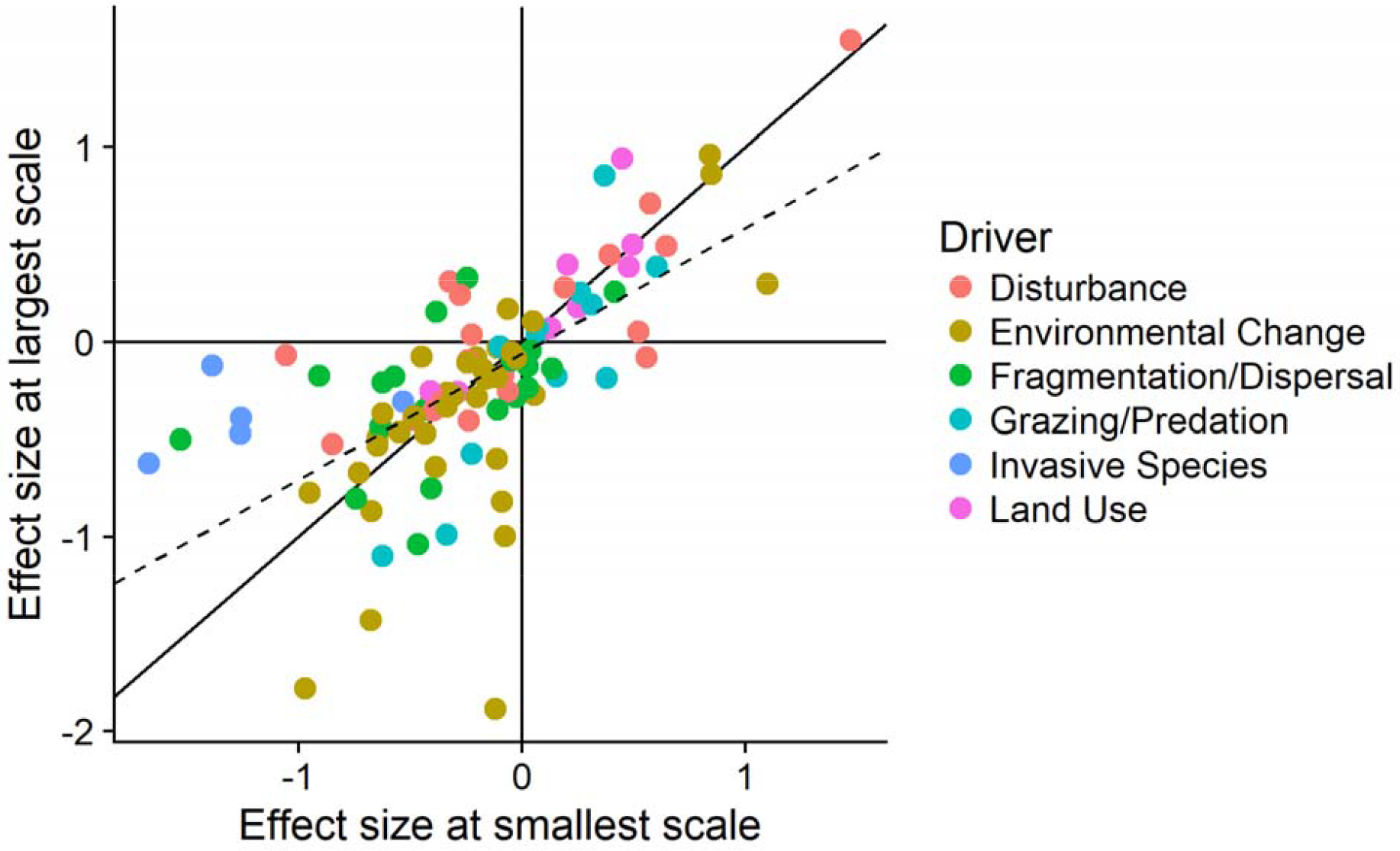
Results of a meta-analysis of scale-dependent responses to a number of different ecological drivers (see Appendix 2). Points represent the log response ratio comparing species richness in control compared to treatments in a given comparison measured at the smallest (x-value) and largest (y-value) scale. The solid line indicates the 1: 1 line expected if effect sizes were not scale-dependent. Points above and below this line indicate effect sizes that are larger or smaller, respectively, as scale increases; points in the upper left and lower right quadrats represent cases where the direction of change shifted from positive to negative, or vice versa, with increasing scale. The dashed line indicates the best fit correlation, which is significantly different than the 1:1 line (P<0.01), indicating that overall, effect sizes tend to be larger at smaller scales than at larger scales. Colors for points indicate categorizations into different ecological drivers.

Given the poor fit of the 1:1 line to predict effect sizes across scales, we can be reasonably certain that scale-dependence is important. However, we can also use the information to evaluate the nature of this scale-dependence in a deeper way. For example, the simple alternative hypotheses we presented in Figure 1 illustrate different ways that scale-dependence can occur. If biodiversity change is ‘well-behaved’ as in Figure 1A, we might expect that there would be a consistent increase in the difference in species richness with increasing scale. This would mean that points should be more likely to fall below the 1:1 line in the negative-negative (lower left) quadrat (larger reductions in richness at larger scales); and above the 1:1 line in the positive-positive (upper right) quadrat (larger increases in richness at larger scales). While there are many cases that fall in these zones, this pattern is in fact less frequent than the opposite, where effect sizes tend to decrease with increasing scale. Indeed, an unconstrained regression shows that the slope is significantly positive, but significantly less than 1 (slope=0.65±0.074; p<0.01). This relationship is still relatively weak, but explains more of the variation than the regression constrained to 1:1 in effect sizes across scales (*R*^2^ = 0.43), and suggests that overall effect sizes at smaller scales are on average larger than effect sizes at larger scales.

We categorized the different ecological drivers from these studies, and found that studies from three drivers in particular— invasive species, land use change, and grazers/predators—showed significantly larger effect sizes at small compared to larger spatial scales (see supplemental Figure S1). These findings reinforce an earlier meta-analysis by Powell et al. (2011), who found that the effect sizes of invasive species on plant richness systematically declined with increasing spatial scale. Because such small-scale estimates of biodiversity change are often used as inputs, for example, into ‘biodiversity scenarios’ models that project future diversity loss (e.g., Alroy 2017, Newbold et al. 2017), we suspect that these analyses may overestimate the actual change observed at larger management-relevant scales.

## Towards a framework to estimate scale-dependent multidimensional biodiversity change

Our above empirical review emphasizes what has been suspected for some time, but not heretofore systematically quantified; comparisons of biodiversity and its change from site to site are not well captured by a single number (Figure 2, 3) nor a single scale of observation (Figure 4). However, from a methodological perspective, the recognition that biodiversity is multidimensional and scale-dependent is nothing new. There have been hundreds of publications aimed at developing, analyzing and comparing various methodologies that can capture some of this complexity.

Three of the most commonly used and recommended approaches for dealing with the multidimensional scaling problem in biodiversity studies include: (1) converting species richness into other measures that incorporate the relative abundances of species, such as Shannon’s or Simpson’s diversity indices (e.g., Hill 1973, Lande 1996, Jost 2006, 2007); (2) comparing species richness values after controlling for sampling effort in the numbers of samples or individuals through rarefaction (e.g., Hurlbert 1971, Simberloff 1972, Gotelli and Colwell 2001, Cayuela et al. 2015); and (3) extrapolating species richness values to a hypothetical asymptote based on the estimation of the number of undetected species (e.g., Chao 1984, Colwell and Coddington 1994, Chao et al. 2009). Importantly, these have been combined into a single statistical framework that uses both interpolation and extrapolation (e.g., Chao and Jost 2012, Colwell et al. 2012), and can be applied to Hill numbers, a family of diversity metrics that give different weighting to the importance of rare and common species (e.g., Chao et al. 2014a).

Despite these advances, this framework does not fully solve the problem of spatial scale. For example, rarefactions that control for the numbers of individuals will still identify scale-dependent species richness rankings that depend on the numbers of individuals to which species richness is rarefied (e.g., Gotelli and Colwell 2001, Cao et al. 2007, Chase and Knight 2013, Cayuela et al. 2015). On the other hand, diversity extrapolations compare only the maximum diversity in a community at a hypothetical sampling asymptote (e.g., the largest scale in Figure 1), which may require different levels of sampling intensity to achieve for different treatment groups (Chao et al. 2009) and do not capture differences in the rise to that asymptote. Metrics that quantify differences in evenness (e.g., Shannon’s, Simpson’s), can capture some of the differences observed (e.g., Hill 1973, Chao et al. 2014a), but as we described above, few studies use more than a single biodiversity metric which is necessary in order to fully evaluate the complexity of biodiversity change, nor do they consider what any similarities or differences in the metrics mean.

## Constructing and comparing elements of rarefaction curves

Gotelli and Colwell (2001) contrasted several types of rarefaction and accumulation curves which contain complementary information on how species diversity varies with sampling effort and scale. Here, we focus on two that are useful for making comparisons across scales and dissecting the relationships between different factors associated with biodiversity measurements: sample-based rarefaction and individual-based rarefaction.

Sample-based rarefaction (SBR) considers increases in the numbers of species as the numbers of sampling units increases (see Chiarucci et al. 2008 for early uses of this concept). The rarefaction curve is an average of different accumulations, in which the ordering of the accumulated samples is random. In our discussion, we modify the original SBR which is not spatially explicit to create the spatial SBR (sSBR), which keeps spatial structure explicit (Figure 5A). That is, one starts with a single plot, then adds the closest plot, etcetera until all plots are included, and the processes are repeated across starting plots (Chiarucci et al. 2009) (we prefer the sSBR terminology rather than ‘spatially constrained rarefaction’ coined by Chiarucci et al. [2009], because their term does not differentiate whether the rarefaction is sample- or individual-based). It is important to note that sSBR is one form of what investigators have generically called species-area relationships (Scheiner 2003). The sSBR, however, differs in some important ways from the nested species area relationships that are most commonly used in macroecological studies (e.g., Storch 2016), even though it is possible to estimate one from the other with certain assumptions (e.g., Azaele et al. 2015, Kunin et al. 2018). We focus here on sSBR curves and their derivatives, rather than nested species area relationships, because the sSBR is more closely aligned with the way that the vast majority of field biologists collect and analyze their data.

**Figure 5.**
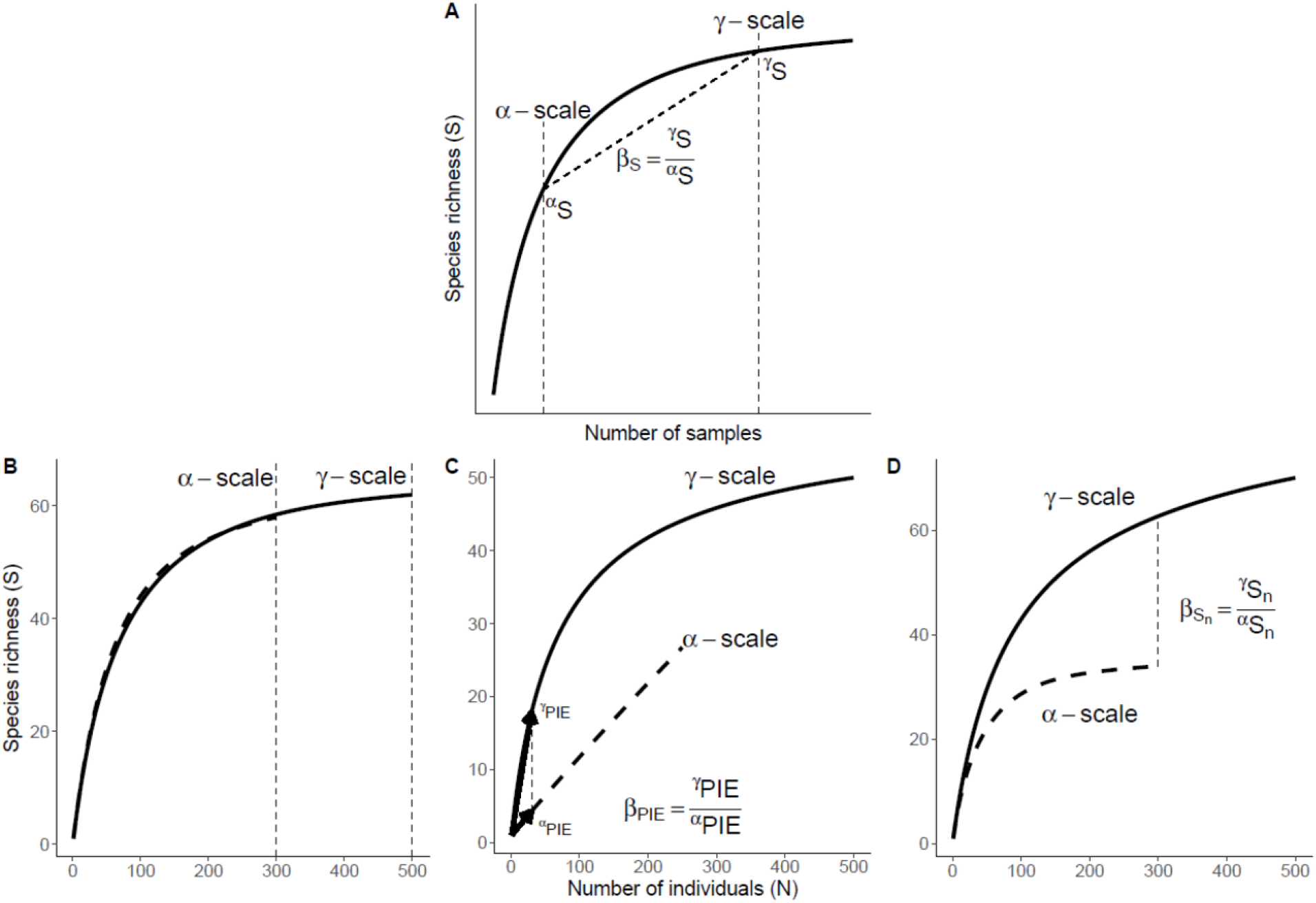
(A) Spatial Sample-Based Rarefaction (sSBR) as defined in text. The average number of species in a sample is defined as the α-scale, and the total number of species across samples is defined as the γ-scale. From this, we can derive local richness (^α^S), regional richness (^γ^S), and β-richness (β_s_=^γ^S/^α^S). (B), (C), (D) individual-based rarefaction curves (IBR) from α-scale samples (dashed lines) and γ-scale samples (solid lines). From this, we can visualize S_n_ and PIE for each scale, and the differences between them (β_Sn_, β_PIE_). Other metrics presented in Table 1 are not shown for clarity. (B) illustrates a case where there is no aggregation in the community, and the IBRs from the α- and γ-scale completely overlap. (C) illustrates a case where there is considerable aggregation in the community, and the IBRs from the α- and γ-scale are completely different; here, β_Sn_ and β_PIE_ are high. (D) illustrates a case where there is some aggregation in the community, it is mostly manifest among the rarer species, such that there is no β_PIE_, but there is β_Sn_.

From the sSBR, we can derive the most commonly used metrics associated with species richness (S) using Whittaker’s (1960) multiplicative diversity partition γ=α*β (Table 1) (see also Jost 2007, Tuomisto 2010) (see Crist and Veech 2006 for an additive partition within a similar approach). Specifically, we can estimate species richness at the small scale, which we call α-diversity and abbreviate as ^α^S. Typically, this will be derived as an average value from a number of replicates or sampling plots. Next, we can estimate species richness at the large scale, which we call γ-diversity and abbreviate as ^γ^S. This will typically be derived from the total numbers of species observed among all replicates or samples (nevertheless, it is important to note that simply adding up species encountered in from multiple sampling plots within a larger regional extent will typically be far from a complete survey of the true γ-diversity in that entire extent). Finally, we can estimate the turnover in species from smaller to larger scales as β-diversity (β=γ/α), which we abbreviate as β_S_.

**Table 1.** Definitions and interpretations of the biodiversity metrics for scale-explicit analyses.

| Metric | Definition | Interpretation |
| --- | --- | --- |
| $N$ | Total numbers of individuals | Measure of density of individuals. Because $N$ scales roughly linearly with area (i.e. density is scale-independent) we do not need to measure at multiple scales. |
| $\alpha_S, \gamma_S$ | Observed richness of species from α-scale (average of observations) and from γ-scale (sum of observations) local scale observations. | Number of species at local scale ($=\alpha$-diversity) and large scale ($=\gamma$-diversity) |
| $\alpha_{S_n}, \gamma_{S_n}$ | The expected richness for $n$ randomly sampled individuals (Hurlbert 1971). Can be calculated from α- or γ-scale. | Estimate of richness at α- or γ-scale after controlling for differences due to aggregation and number of individuals (i.e., only reflects SAD) |
| $\alpha_{PIE}, \gamma_{PIE}$ | Probability of intraspecific encounter ($S_{n=2} - S_{n=1}$, Hurlbert 1971, Olszewski 2004) | Measure of evenness, slope at base of the rarefaction curve, and sensitive to common species |
| $\alpha_{S_{PIE}}, \gamma_{S_{PIE}}$ | Equally abundant species needed to yield PIE (Jost 2006) ($=1/(1-PIE)$) | Effective number of species of PIE ($=1$-Simpson's) that is easier to compare with $S$ |
| $\beta_S$ | Ratio of total treatment $\gamma_S$ and average plot $\alpha_S$ (Whittaker, 1960) | More species turnover results in larger $\beta_S$ because of increases in spatial aggregation, $N$, and/or unevenness of the SAD. |
| $\beta_{PIE}$ | Ratio of total treatment $\gamma_{PIE}$ and $\alpha_{PIE}$ (Olszewski 2004) | Higher difference between $\gamma_{PIE}$ and $\alpha_{PIE}$ indicates higher levels of aggregation. |
| $\beta_{S_n}$ | Ratio of total treatment $\gamma_{S_n}$ and $\alpha_{S_n}$ | Like $\beta_S$ but emphasizes aggregation due to common and rare species. |
| $\beta_{S_{PIE}}$ | Ratio of total treatment $\gamma_{S_{PIE}}$ and $\alpha_{S_{PIE}}$ (Jost 2007) | Like $\beta_S$ but emphasizes aggregation due to common species. |

Our definitions of α-, β- and γ-diversity are not intended to define any particular features of a given community, such as zones where species interact and coexist, or where a given species pool is defined. It is our view that it is unlikely that any discrete scale can be reasonably defined to be explicitly ‘local’ or ‘regional’, but rather these demarcations are a somewhat arbitrary way to define smaller and larger scale estimates of diversity and their comparison. Nevertheless, for specific comparisons, it is critical to make sure that α- and γ-scale comparisons are done with similar sampling effort (e.g., Gotelli and Colwell 2001). We also note that our use of β-diversity in the context of the diversity partition is intended to capture variation within a given treatment of an ecological driver, and is quite different than approaches that compare compositional differences (Tuomisto 2010, Anderson et al. 2011).

Critically, the spatially explicit and sample-based structure of the sSBR and values derived from it (e.g., α–, β–, γ–diversity) are jointly determined by the total numbers of individuals in a community (N), the shape of the SAD (including total species numbers and their relative abundances), and their intraspecific aggregations or clumping, and these components can have bi-directional effects. Furthermore, changes in any one of the components can lead to similar looking sSBRs, and changes in two or more components can lead to quite complex outcomes (e.g., Chase and Knight 2013, Powell et al. 2013). Thus, one of our main goals in evaluating the multidimensional and scale-dependent way by which biodiversity changes across space or time is to dissect the influence of each of the components on the observed sSBR. While we discuss N, the SAD and aggregation as the components influencing changes in diversity, it is important to note that our approach is agnostic to mechanism. Thus, while changes in observed S that are correlated with changes in total N in the community is consistent with some forms of the ‘more individuals hypothesis’ (sensu Srivastava and Lawton 1999), for example, we cannot necessarily infer that change in N causes change in S in a unidirectional way, but simply that they are correlated (e.g., Storch et al. 2018).

Whereas the sSBR is the most complex curve that preserves information on differences among two or more communities N, SAD and aggregation, the individual-based rarefaction (IBR) is the most simple (Figure 5B). Specifically, in the IBR, individuals are pooled across all samples in a treatment and selected at random, so that any spatial structure in intraspecific aggregation and differences in the density of individuals in sampling plots is removed (e.g., Hurlbert 1971, Simberloff 1972, Gotelli and Colwell 2001). While the distance along the x-axis (number of individuals) is determined by N, the shape of the IBR is determined only by the SAD. The height of the IBR is the total numbers of species (S) observed among all of the samples pooled, and the steepness of the slope near the origin is determined by the relative abundances (evenness) of species represented in the pooled samples (Hurlbert 1971, Gotelli and Colwell 2001, Olszwewski 2004, Chase and Knight 2013, Cayuela et al. 2015).

From the IBR, we can derive a number of parameters of interest (Figure 5B-D; Table 1). To derive these parameters across scales, we calculate two IBRs, one using only the N and S from samples collected at the α-scale (i.e., from a single replicate or plot, or several plots within a locality embedded within a metacommunity) and one from the γ-scale across all replicates or localities within a given treatment or type (a multi-scale version of this approach, including benchmark tests and evaluation of statistical error, is presented in McGlinn et al. 2018). From the α- and γ-scale IBRs, and the measures we can derive from these, we can begin to understand the relationship between N, the SAD and aggregation and the observed diversity patterns.

### Disentangling the influence of N by comparing rarefied richness values

We can first examine whether any differences in ^α^S or ^γ^S observed from the sSBR result from differences in total N among treatments. Specifically, with the IBR, we can calculate the numbers of species expected when rarefied to a common N, known as rarefied richness, abbreviated as S_n_. Because differences in rarefied richness are also scale-dependent (e.g., Cao et al. 2007, Chase and Knight 2013, Cayuela et al. 2015), we advise comparing S_n_ at multiple values of N. Further, we can compare values of S_n_ at different scales of sampling, which we denote ^α^S_n_ and ^γ^S_n_, respectively.

### Disentangling the influence of SAD by comparing elements of individual-based rarefaction curves

If the shapes of the IBRs differ between treatments, we can infer that SADs differ via changes in the numbers of species (S), the relative abundances (evenness) of species, or both (as shown above in Figure 1). To evaluate this, we can perform statistical comparisons at two extremes of the IBR, the upper limit (S) at the right extreme of the curve and the slope at the base of the IBR. The upper limit of the IBR can be calculated in a number of ways, depending on which sampling controls are most appropriate, including the rarefied richness S_n_ at standardized values of N (e.g., Gotelli and Colwell 2001), S calculated to a fixed coverage of sample completeness (e.g., Chao and Jost 2012) or an extrapolation to a hypothetical asymptote where coverage is complete (e.g., Chao 1984, Chao et al. 2009, 2014a). The slope at the base of the IBR, as described above, is equivalent to the bias-corrected probability of interspecific encounter (PIE) (Hurlbert 1971, Olszwewski 2004), and represents a measure of evenness in the community. The PIE can be measured from samples at the α-scale (^α^PIE), or at the γ-scale (^γ^PIE). Because PIE is the same as 1-Simpson’s diversity index, we can convert it to an effective number of species (S_PIE_) as above, and calculate this quantity at each spatial scale: ^α^S_PIE_ and ^γ^S_PIE_. Both PIE and S_PIE_ can provide useful information for comparisons between two or more communities (e.g., Dauby and Hardy 2012). Because S and PIE (or S_PIE_) are indicative of different parts of the IBR, and also reflect different locations along the continuum of the Hill numbers (e.g., Hill 1973, Jost 2006), they conceptually link the IBR with Hill numbers, which differentially weight common versus rare species.

### Disentangling β-diversity due to aggregation using individual-based rarefaction curves

Raw measures of β-diversity from the sSBR are influenced by N and the shape of the SAD, as well as by spatial aggregation. To tease apart the specific influence of aggregation on β-diversity, we can compare the γ-scale IBR (which randomly samples individuals regardless of their spatial position), to the α-scale IBR (which preserves effects of aggregation within plots or replicates).

First, we can compute β_PIE_ as difference between the slopes defined by ^α^PIE and ^γ^PIE at the base of the respective IBRs (Figure 5B-D) (Olszwewski 2004, Dauby and Hardy 2012). And we can convert this to an effective number of species by taking the ratio of small-scale (^α^S_PIE_) to large-scale (^γ^S_PIE_) to yield *β*_*S*_PIE__ (Jost 2007, Tuomisto 2010). Second, we can compute *β*_*S*_n__ as the ratio of the number of species expected from a given *n* from the γ-scale IBR (^γ^S_n_) to the number of species observed from a given *n* from the α-scale IBR (^α^S_n_). Three qualitative outcomes of these different components of β-diversity are possible. If there is no aggregation, the γ-scale and α-scale IBRs will not differ in their shapes and there will be no differences due to spatial clumping in the landscape. Thus β_PIE_, *β*_*S*_PIE__, and *β*_*S*_n__, will equal 1 (Figure 5B). Second, if there is strong aggregation in the community, there can be large differences in γ-scale and α-scale IBRs, leading to high levels of β_PIE_ (and *β*_*S*_PIE__), as well as *β*_*S*_n__ (Figure 5C). Finally, it is possible that the γ-scale and α-scale IBRs can differ in their shapes, but do not differ in PIE, which measures turnover in the most common species, but misses changes that might be due to aggregation of rarer species in the community. Here, we would observe β_PIE_ (or *β*_*S*_PIE__) equal to 0, but *β*_*S*_n__ greater than 0 (Figure 5D).

### Recommended protocol for analyses dissecting biodiversity data with a case study

Next, we provide a recipe for using the multiple metrics at multiple scales, as presented above and overviewed in Table 1. We describe how one might address these questions generically, and illustrate this recipe with a re-analysis of an experiment on the effect of nutrient additions on macroinvertebrates and amphibians in experimental ponds, showing how deeper insights can be gained than are possible with a more traditional perspective (for full details, see Chase 2010).

R code for these analyses, as well as a series of vignettes and another case study, are available in *mobr* statistical package (https://github.com/MoBiodiv/mobr) (McGlinn et al. 2018). We specifically used one-way PERMANOVA to assess the treatment effect on estimates of diversity measured at the α-scale (=1 mesocosm) (treatment group labels were permuted 999 times to generate a null distribution, and then the tail probability for a treatment effect was evaluated by comparing the observed difference to the null distribution). At the γ-scale (=15 mesocosms), the null distribution was generated by permuting treatment group labels across samples, pooling the groups, and then calculating the difference in diversity between treatments for each permutation. Results are illustrated in Figure 6; some results are not shown for brevity.

**Figure 6.**
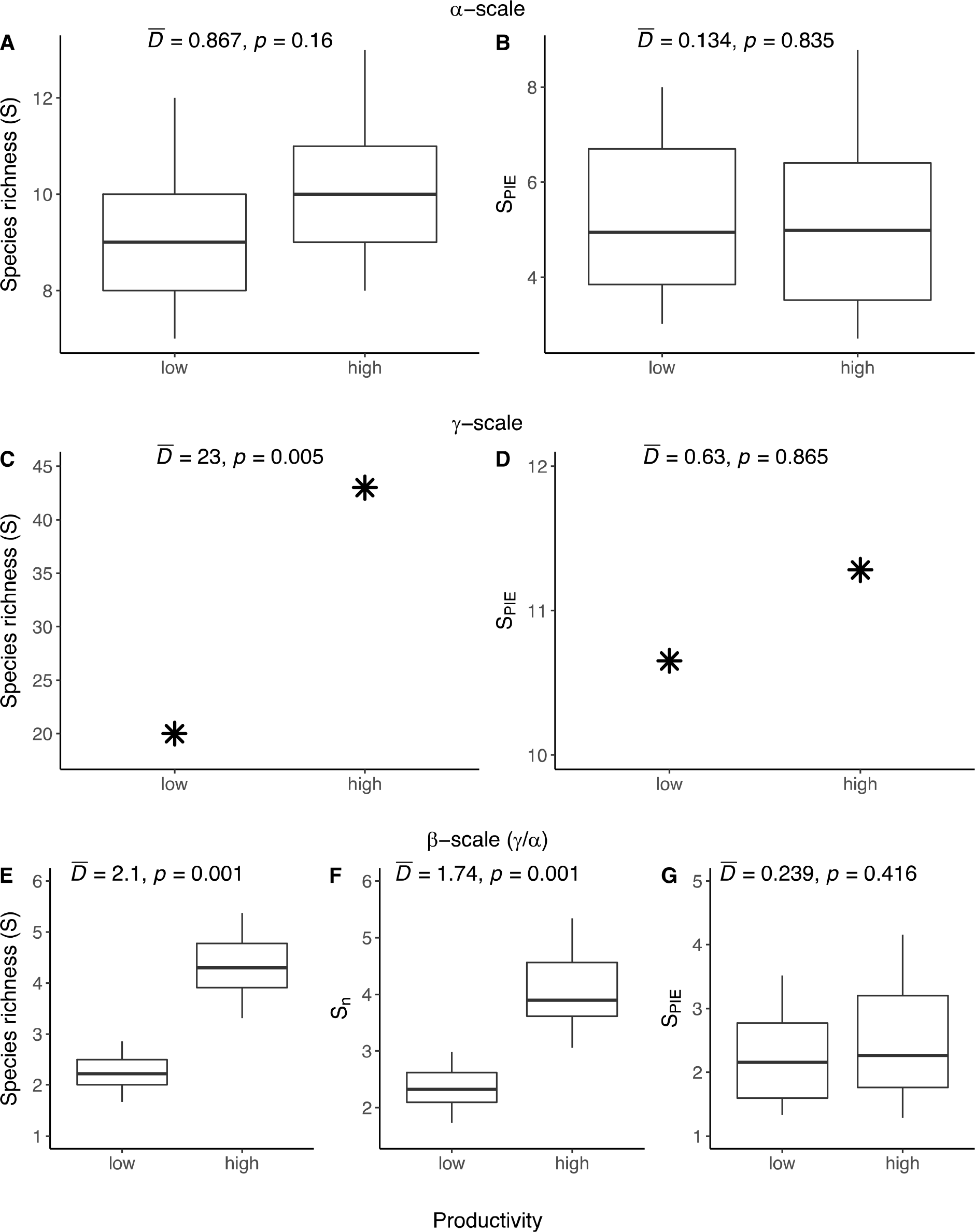
Effect of nutrient additions on several measurements of biodiversity from Table 1 (see data in Appendix 3). Each biodiversity measure was calculated at the α-scale (1 mesocosm) (Panels A,B), γ-scale (15 mesocosms) (Panels C,D), as well as the β-scale (i.e. turnover across scales, γ/α)(Panels D,E,F). See Chase (2010) for details on the experimental design and the mobr package (McGlinn et al. submitted, https://github.com/MoBiodiv/mobr) for details on the statistical methods

#### Step 1: Do treatments affect the numbers of individuals of all species (N)?

If treatments differ in N, then any difference in S between treatments may, in part, be due to treatment effects on N. We expect that N scales linearly with increasing sampling (unlike S), and so it should not matter whether the average N per plot (α-scale), or the total N across plots (γ-scale), are compared. In our case study, we found no influence of the nutrient addition treatment on N measured at either scale (P>0.3; not shown in Figure).

#### Step 2: Do treatments affect diversity responses at the α-scale?

Here, we can compare several types of α-scale diversity. Comparing effects of the treatment on *^α^S* with treatment effects on *^α^S*_n_ allow us to discern any influence of changes in N on the results. For example, if any treatment-level effects on *^α^S* disappear for *^α^S*_n_, we can conclude that the effect was due to treatment effects on N. However, if there remains a difference in *^α^S*_n_, we can conclude that the treatment effect on *^α^S* was due, at least in part, to changes in the SAD. It is even possible that the effects can shift between *^α^S* and *^α^S*_n_. For example, McCabe and Gotelli (2000) showed that *^α^S* was higher for stream invertebrates in undisturbed treatments, but that this effect was reversed when the effect of disturbance on N was discounted; *^α^S*_n_ was higher in disturbed treatments. A difference in *^α^S*_PIE_ implies a change in the dominance patterns of common species in a community. Alternatively, it is possible that *^α^S*_PIE_ could not vary (or vary little), while there is still an influence of the treatment on *^α^S*_n_. This would imply that rarer species are responding to the treatment.

From our case study, we observed that *^α^S* did not differ between the treatments (Fig. 6A), and because there was no difference in N, the effects of N on *^α^S* and *^α^S*_n_ did not differ among treatments (result not shown in figure). Furthermore, there was no influence of treatment on *^α^S*_PIE_ (Figure 6B), indicating that nutrient addition treatment had no identifiable effects on N or any measures of diversity that capture differences in the shape of the SAD at the local scale. Had we stopped here, we might conclude that there was no interesting influence of nutrients on biodiversity. However, by looking at the γ-scale, we see that this conclusion is wrong.

#### Step 3: Do treatments affect diversity responses at the γ-scale?

Comparisons of γ-scale diversity measures are a straightforward extension of the α-scale measures, except we use the totals observed in all samples, rather than the average number from a single sample. And the relationships among the measures are the same, where *^γ^S* is the total richness of species, *^γ^S*_n_ is the richness of species rarefied to a common N, and *^γ^*PIE and *^γ^S*_PIE_ consider the relative abundances of species and conversion to an effective number of species, respectively.

From our case study, and in contrast to the local-scale result, we found a significant increase in *^γ^S* (Fig. 6C) and *^γ^S*_n_ (not shown) with nutrient additions. However, there was no influence of nutrient additions on *^γ^PIE* (not shown) or *^γ^S*_PIE_ (Figure 6D). This suggests that it is the rare species end of the SAD that changed among the treatments; nutrient enrichment allowed more rare species to persist regionally, but not locally. Further, the qualitatively different results at the γ-and α-scale imply that β-diversity was influenced by the nutrient addition treatment.

#### Step 4: Do treatments affect β-diversity?

First, we can test for differences in *β_S_* among treatments. However, as discussed above, any differences could occur due to a combination of changes in N, the SAD or aggregation. If the treatment influences aggregation, we would expect a difference in *β*_*S*_n__ between treatments. If the treatment affects the aggregation of common species, we would also expect to see treatment-level effects on *β*_*S*_PIE__.

As anticipated from the scale-dependent results we observed at the α– and γ-scale (Figure 6A, 6C), we found strong effects of nutrient addition on *β*_*S*_ (Fig. 6E), and on *β*_*S*_n__ (Fig. 6F). And because we found no difference in the α– and γ-scale results for S_PIE_, we unsurprisingly found no treatment-level effect on *β*_*S*_PIE__ (Fig. 6G). Overall, these patterns suggest that a core group of common species were present across replicates in both treatments, and that a group of rarer species were largely responsible for the treatment-level responses at the γ-scale. Specifically, there was more turnover among those rare species in the high nutrient treatment, which may have resulted from ecological drift and/or priority effects.

## Conclusions

There are multiple pathways by which ecological factors can influence biodiversity across multiple scales, but most studies continue to rely on comparisons of a single summary variable—usually species richness (S)—at a single spatial scale. As a consequence, despite thousands of published studies quantifying how species richness changes in response to natural and anthropogenic drivers, we know much less than we think about how and why biodiversity changes from place to place and time to time. This is particularly problematic when trying to achieve synthesis across multiple studies of multiple ecological drivers, through meta-analyses and other means, because effect sizes are highly confounded by spatial scale (see also Chase and Knight 2013). We are currently limited in our ability to create realistic ‘biodiversity scenarios’ models that project future biodiversity loss in response to changing ecological conditions.

To move forward, it is critical to consider the multidimensional and scale-dependent nature of biodiversity and its change. Fortunately, there is a rich literature on other measures of biodiversity that can explicitly complement comparisons of S, and there are easy ways that empiricists can explicitly deal with issues of spatial scaling (i.e., replicates nested within treatments). We have advocated for measures of biodiversity and its scaling that can explicitly be used to disentangle the influence of N, the SAD and aggregation via consideration of different aspects of the individual-level rarefaction curve that emphasize different underlying components (e.g., S_n_, S_PIE_, at the α and γ scales). For a majority of studies, these can be estimated in a straightforward way with data that are already, or could be, collected.

The approach we advocate will often require more complex collecting and reporting of data (i.e., absolute species abundances at multiple scales) and more complex analyses of multiple response variables. This will create a more nuanced view of what biodiversity is and how it varies—biodiversity is not a single number, and it cannot be compared at a single scale to estimate how it responds to ecological factors. However, this more nuanced view is necessary to resolve long-standing debates. For example, Blowes et al. (2017) recently used this approach to show that the debate about whether environmental versus historical biogeographic controls influence global biodiversity patterns of reef-associated fishes can be progressed by dissecting patterns of species richness in a scale-explicit way.

Finally, there are many extensions of the approach that we advocate which are necessary to be able to fully understand, and synthesize, how biodiversity changes in time and space. First, as we mentioned above, the approach we have taken here views scale in a discrete two-scale way (e.g., α, β, γ–diversity), while scale is continuous. In a companion paper, we develop a multiscale methodology (McGlinn et al. 2018). Second, we have only focused on taxonomic diversity, although interest in other measures of diversity has increased greatly in recent years. These measures, such as functional and phylogenetic diversity, show patterns of scaling similar to taxonomic diversity (e.g., Morlon et al. 2011, Smith et al. 2013) and will thus show scale-dependence when making comparisons. Approaches similar to those advocated here are emerging for these other types of diversity (Chao et al. 2014b, 2015, Chiu et al. 2104), and so we anticipate that a family of approaches for comparing scale-dependent diversity responses to ecological drivers at multiple levels of organization will soon emerge.

## Acknowledgements

This paper emerged from several working group meetings funded by the German Centre of Integrative Biodiversity Research (iDiv) Halle-Jena-Leipzig (funded by the German Research Foundation; FZT 118) and the Alexander von Humboldt Foundation as part of the Alexander von Humboldt Professorship of TMK. DJM was also supported by the College of Charleston startup funding. NJG was supported by U.S. NSF DEB 1257625. BJM was supported by USDA Hatch grant to MAFES #1011538 and NSF ABI grant #1660000. We thank J. Belmaker, N. Sanders, D. Storch, two anonymous reviewers and the handling editor, and a number of other colleagues that helped us to develop and reform these ideas and tools.

## Supplementary Information

**Figure S1.**
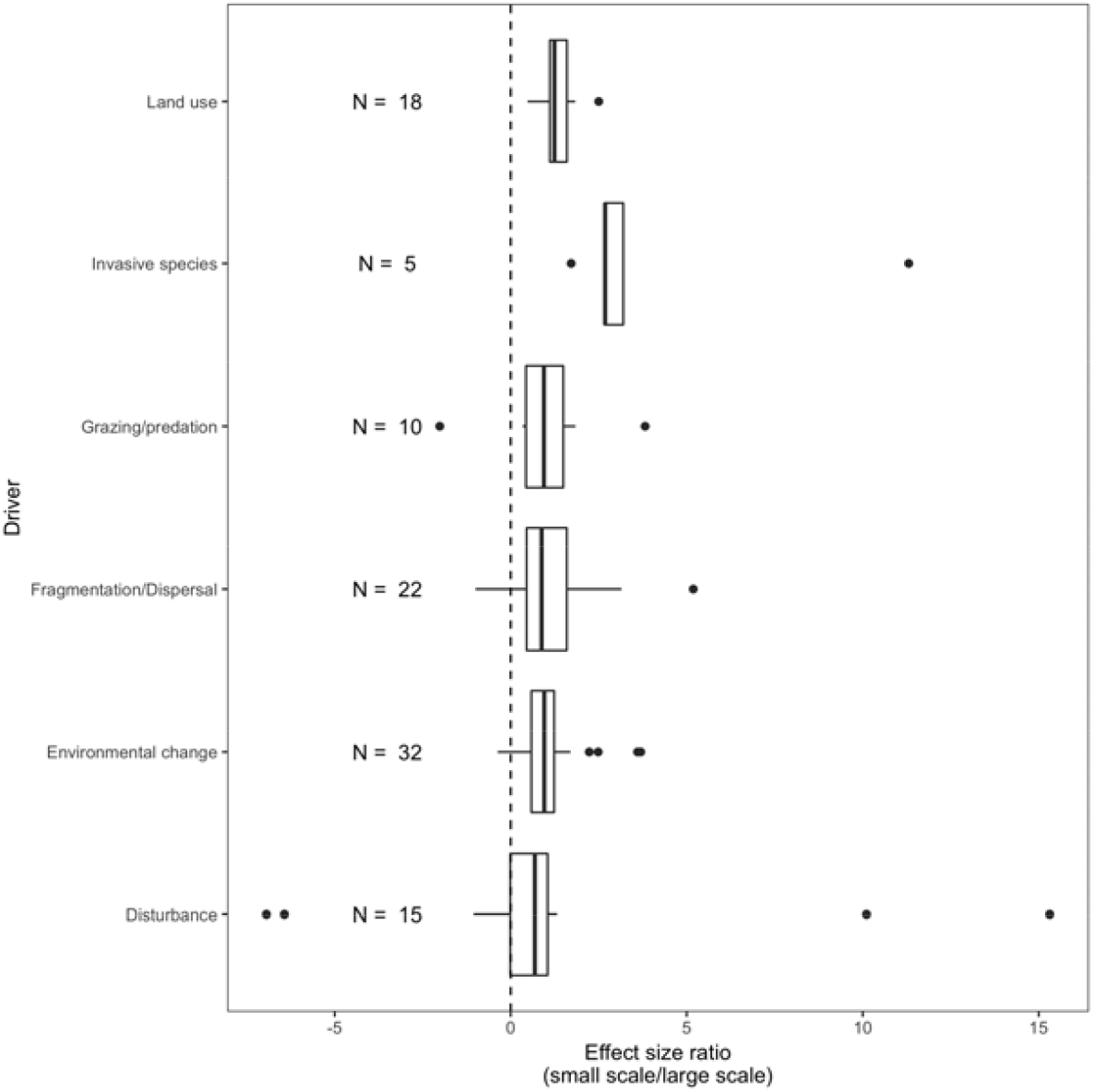
Results showing the ratio of log-response ratio effect sizes from experiments where species richness responses were measured at two spatial scales (small scale/large scale). The dashed line at 0 would indicate studies where the effect sizes were the same at the smaller and larger scale. For each category of ecological driver, the means are above 1, indicating that the measured effect size is larger at the smaller relative to larger size, and this difference is statistically significant for land use, invasive species, and grazing/predation.

## Appendix 1

Raw data and references for rarefaction curves and analyses presented in Figures 2 and 3. Subset of studies presented in McGill 2011b.

File available here: https://www.dropbox.com/sh/s3vmrwya1b7khz2/AAAnWDucBJYdxPdT_tOk2olOa?dl=Q (will be deposited in appropriate repository on acceptance)

## Appendix 2

Effect sizes, metadata, and references for studies used in the scale-dependent metaanalysis presented in Figure 4.

File available here: https://www.dropbox.com/sh/s3vmrwya1b7khz2/AAAnWDucBJYdxPdT_tQk2olQa?dl=0 (will be deposited in appropriate repository on acceptance)

## Appendix 3

Raw data for the analyses of invertebrate and amphibian communities in nutrient addition and reference treatments presented in Figure 6. From Chase (2010).

File available here: https://www.dropbox.com/sh/s3vmrwya1b7khz2/AAAnWDucBJYdxPdT_tOk2olOa?dl=0 (will be deposited in appropriate repository on acceptance)

## Literature Cited

Alroy, J., (2017). Effects of habitat disturbance on tropical forest biodiversity. Proc. Natl. Acad. Sci. 114, 6056–6061.

Azaele, S., Maritan, A., Cornell, S.J., Suweis, S., Banavar, J.R., Gabriel, D. and Kunin, W.E., (2015). Towards a unified descriptive theory for spatial ecology: Predicting biodiversity patterns across spatial scales. Methods Ecol. Evol. 6, 324–332.

Anderson, M.J., Crist, T.O., Chase, J.M., Vellend, M., Inouye, B.D., Freestone, A.L., et al, (2011). Navigating the multiple meanings of β diversity: a roadmap for the practicing ecologist. Ecol. Lett., 14, 19–28.

Andrew, M.E., Wulder, M.A., Coops, N.C. and Baillargeon, G., (2012). Beta-diversity gradients of butterflies along productivity axes. Global Ecol. Biogeog. 21: 352–364.

Baselga, A., (2010). Partitioning the turnover and nestedness components of beta diversity. Global Ecol. Biogeog., 19, 134–143.

Blowes, S. A., Belmaker, J., and Chase, J. M. (2017). Global reef fish richness gradients emerge from divergent and scale-dependent component changes. Proc. R. Soc. B 284, 20170947

Bürkner, P-C. (2017). brms: An R package for Bayesian multilevel models using Stan. J. Stat. Soft., 80, 1–28.

Cao, Y., Hawkins, C.P., Larsen, D.P. and Van Sickle, J., (2007). Effects of sample standardization on mean species detectabilities and estimates of relative differences in species richness among assemblages. Am. Nat., 170, 381–395.

Cayuela, L., Gotelli, N.J. and Colwell, R.K., (2015). Ecological and biogeographic null hypotheses for comparing rarefaction curves. Ecol. Monog., 85, 437–455.

Chao, A., (1984). Nonparametric estimation of the number of classes in a population. Scand. J. Stat., 11, 265–270

Chao, A. and Jost, L., (2012). Coverage-based rarefaction and extrapolation: standardizing samples by completeness rather than size. Ecology, 93, 2533–2547.

Chao, A., Colwell, R.K., Lin, C.W. and Gotelli, N.J., (2009). Sufficient sampling for asymptotic minimum species richness estimators. Ecology, 90, 1125–1133.

Chao, A., Gotelli, N.J., Hsieh, T.C., Sander, E.L., Ma, K.H., Colwell, R.K. and Ellison, A.M., (2014a). Rarefaction and extrapolation with Hill numbers: a framework for sampling and estimation in species diversity studies. Ecol. Monog., 84, 45–67.

Chao, A., Chiu, C.H. and Jost, L., (2014b). Unifying species diversity, phylogenetic diversity, functional diversity, and related similarity and differentiation measures through Hill numbers. Annu. Rev. Ecol. Syst, 45, 297–324.

Chao, A., Chiu, C.H., Hsieh, T.C., Davis, T., Nipperess, D.A. and Faith, D.P., (2015). Rarefaction and extrapolation of phylogenetic diversity. Methods Ecol. Evol., 6, 380–388.

Chase, J.M. and Leibold, M.A., (2002). Spatial scale dictates the productivity–biodiversity relationship. Nature, 416, 427–430.

Chase, J.M., (2010). Stochastic community assembly causes higher biodiversity in more productive environments. Science, 328, 1388–1391.

Chase, J. M., and Knight, T. M. (2013). Scale-dependent effect sizes of ecological drivers on biodiversity: why standardised sampling is not enough. Ecol. Lett., 16, 17–26.

Chiarucci, A., Bacaro, G., Rocchini, D. and Fattorini, L., (2008). Discovering and rediscovering the sample-based rarefaction formula in the ecological literature. Comm. Ecol., 9, 121–123.

Chiarucci, A., Bacaro, G., Rocchini, D., Ricotta, C., Palmer, M. and Scheiner, S., (2009). Spatially constrained rarefaction: incorporating the autocorrelated structure of biological communities into sample-based rarefaction. Comm. Ecol., 10, 209–214.

Chiu, C.H., Jost, L. and Chao, A., (2014). Phylogenetic beta diversity, similarity, and differentiation measures based on Hill numbers. Ecol. Monog., 84, 21–44.

Colwell, R.K. and Coddington, J.A., (1994). Estimating terrestrial biodiversity through extrapolation. Phil. Trans. Roy. Soc B, 345, 101–118.

Colwell, R.K., Chao, A., Gotelli, N.J., Lin, S.Y., Mao, C.X., Chazdon, R.L. et al. (2012). Models and estimators linking individual-based and sample-based rarefaction, extrapolation and comparison of assemblages. J. Plant Ecol., 5, 3–21.

Condit R, Lao, S, Pérez, R. Dolins, S. B., Foster, R. B. and Hubbell, S. P. (2012). Barro Colorado Forest Census Plot Data, 2012 Version. http://dx.doi.org/10.5479/data.bci.20130603

Dornelas, M., Gotelli, N.J., McGill, B., Shimadzu, H., Moyes, F., Sievers, C. and Magurran, A.E., (2014). Assemblage time series reveal biodiversity change but not systematic loss. Science, 344, 296–299.

Dauby, G. and Hardy, O.J., (2012). Sample-based estimation of diversity sensu stricto by transforming Hurlbert diversities into effective number of species. Ecography, 35, 661–672.

Dumbrell, A.J., Clark, E.J., Frost, G.A., Randell, T.E., Pitchford, J.W. and Hill, J.K., (2008). Changes in species diversity following habitat disturbance are dependent on spatial scale: theoretical and empirical evidence. J. Appl. Ecol., 45, 1531–1539.

Gardezi, T. and Gonzalez, A. (2008). Scale dependence of species-energy relationships: Evidence from fishes in thousands of lakes. Am. Nat. 171, 800–815.

Gelman, A., Goodrich, B., Gabry, J., Ali, I. (2017) R-squared for Bayesian regression models. http://www.stat.columbia.edu/~gelman/research/unpublished/

Gotelli, N.J. and Colwell, R.K., (2001). Quantifying biodiversity: procedures and pitfalls in the measurement and comparison of species richness. Ecol. Lett., 4, 379–391.

Harrod, J. (1964). The distribution of invertebrates on submerged aquatic plants in a chalk stream. J. Anim. Ecol., 33, 335–348.

He, F. and Legendre, P., (2002). Species diversity patterns derived from species–area models. Ecology, 83, 1185–1198.

Hill, M.O., (1973). Diversity and evenness: a unifying notation and its consequences. Ecology, 54, 427–432.

Hill, J.K. and Hamer, K.C., (2004). Determining impacts of habitat modification on diversity of tropical forest fauna: the importance of spatial scale. J. Appl. Ecol., 41, 744–754.

Hillebrand, H., Blasius, B., Borer, E.T., Chase, J.M., Downing, J.A., Eriksson, et al., 2018. Biodiversity change is uncoupled from species richness trends: Consequences for conservation and monitoring. Journal Appl. Ecol., 55, 169–184.

Hurlbert, S.H., (1971). The nonconcept of species diversity: a critique and alternative parameters. Ecology, 52, 577–586.

Jost, L., (2006). Entropy and diversity. Oikos, 113, 363–375.

Jost, L., (2007). Partitioning diversity into independent alpha and beta components. Ecology, 88, 2427–2439.

Keil, P., Biesmeijer, J.C., Barendregt, A., Reemer, M. and Kunin, W.E., (2011). Biodiversity change is scale-dependent: an example from Dutch and UK hoverflies (Diptera, Syrphidae). Ecography, 34, 392–401.

Kunin, W.E., Harte, J., He, F., Hui, C., Jobe, R.T., Ostling, A., Polce, C., Šizling, A., Smith, A.B., Smith, K. and Smart, S.M., (2018). Upscaling biodiversity: estimating the species–area relationship from small samples. Ecological Monographs 88, 170–187.

Lande, R., (1996). Statistics and partitioning of species diversity, and similarity among multiple communities. Oikos, 5–13.

Lande, R., DeVries, P.J. and Walla, T.R., (2000). When species accumulation curves intersect: implications for ranking diversity using small samples. Oikos, 89, 601–605.

Legendre, P., 2014. Interpreting the replacement and richness difference components of beta diversity. Global Ecol. Biogeog., 23, 1324–1334.

Livne-Luzon, S., Ovadia, O., Weber, G., Avidan, Y., Migael, H., et al. (2017). Small-scale spatial variability in the distribution of ectomycorrhizal fungi affects plant performance and fungal diversity. Ecol. Lett., 20, 1192–1202.

MacArthur, R.H. and MacArthur, J.W., (1961). On bird species diversity. Ecology, 42, 594–598.

Magurran, A. E., and McGill, B. J. (eds) (2011) Biological diversity: Frontiers in measurement and assessment. Oxford University Press.

May, F., Gerstner, K. . McGlinn, D. J., Xiao, X and Chase, J. M. (2017) mobsim: An R package for the simulation and measurement of biodiversity across spatial scales. bioRxiv (2017): 209502.

Martinson, V.G., Douglas, A.E. and Jaenike, J., (2017). Community structure of the gut microbiota in sympatric species of wild Drosophila. Ecol. Lett., 20, 629–639.

McCabe, D.J. and Gotelli, N.J., (2000). Effects of disturbance frequency, intensity, and area on assemblages of stream macroinvertebrates. Oecologia, 124, 270–279.

McGill, B. J. (2011a). Linking biodiversity patterns by autocorrelated random sampling. Am. J. Bot., 98, 481–502.

McGill, B. J. (2011b). Species abundance distributions. In. A. Magurran and B. McGill (eds) Biological diversity: frontiers in measurement and assessment. Oxford University Press, Oxford (201:1) 105–122.

McGlinn, D.J. and Palmer, M.W., (2009). Modeling the sampling effect in the species–time–area relationship. Ecology, 90, 836–846.

McGlinn, D. Xiao, X., May, F. Gotelli, N. J., Blowes, S. A., Knight, T. M. et al. Submitted. MoB (Measurement of Biodiversity): a method to separate the scale-dependent effects of species abundance distribution, density, and aggregation on diversity change. Submitted to Meth. Ecol. Evol. (available also on https://www.biorxiv.org/content/early/2018/01/07/244103)

Mendes, R.S., Evangelista, L.R., Thomaz, S.M., Agostinho, A.A. and Gomes, L.C., (2008). A unified index to measure ecological diversity and species rarity. Ecography, 31, 450–456.

Morlon, H., Schwik, D. W. Bryant, L. A., Marquet, P. A., Rebelo, A. G. et al. (2011). Spatial patterns of phylogenetic diversity. Ecol. Lett. 14, 141–149.

Newbold, T., Boakes, E.H., Hill, S.L., Harfoot, M.B. and Collen, B., (2017). The present and future effects of land use on ecological assemblages in tropical grasslands and savannas in Africa. Oikos, 126, 1760–1769.

Olszewski, T.D., (2004). A unified mathematical framework for the measurement of richness and evenness within and among multiple communities. Oikos, 104, 377–387.

Palmer, M.W. and White, P.S., (1994). Scale dependence and the species-area relationship. Am. Nat., 144, 717–740.

Pardieck, K. L., Ziolkowski Jr, D. J., Lutmerding, M, Campbell, K., and Hudson, M-. A. R.. 2017. North American Breeding Bird Survey Dataset 1966 - 2016, version 2016.0. U.S. Geological Survey, Patuxent Wildlife Research Center. <www.pwrc.usgs.gov/bbs/rawdata/>; doi:10.5066/F7W0944J.

Powell, K.I., Chase, J.M. and Knight, T.M., (2011). A synthesis of plant invasion effects on biodiversity across spatial scales. Am. J. Bot., 98, 539–548.

Powell, K.I., Chase, J.M. and Knight, T.M., (2013). Invasive plants have scale-dependent effects on diversity by altering species-area relationships. Science, 339(6117), pp.316–318.

Preston, F.W., (1960). Time and space and the variation of species. Ecology, 41, 611–627.

Rahbek, C., (2005). The role of spatial scale and the perception of large‐scale species‐richness patterns. Ecology Letters, 8, 224–239.

R Core Team (2017). R: A language and environment for statistical computing. R Foundation for Statistical Computing, Vienna, Austria. URL http://www.R-project.org/.

Sandel, B. and Smith, A.B., (2009). Scale as a lurking factor: incorporating scale-dependence in experimental ecology. Oikos, 118, 1284–1291.

Scheiner, S.M., (2003). Six types of species-area curves. Global Ecol. Biogeog., 12, 441–447.

Scheiner, S. M., Cox, S.B., Mittelbach, G.G., Osenberg, C. and Kaspari, M., (2000). Species richness, species–area curves and Simpson’s paradox. Evol. Ecol. Res. 2, 791–802.

Simberloff, D., (1972). Properties of the rarefaction diversity measurement. Am. Nat. 106, 414–418.

Smith, A.B., Sandel, B., Kraft, N.J. and Carey, S., (2013). Characterizing scale-dependent community assembly using the functional-diversity-area relationship. Ecology, 94, 2392–2402.

Solar, R.R.D.C., Barlow, J., Ferreira, J., Berenguer, E., Lees, A.C., et al., (2015). How pervasive is biotic homogenization in human-modified tropical forest landscapes? Ecol. Lett., 18, 1108–1118.

Srivastava, D.S. and Lawton, J.H., (1998). Why more productive sites have more species: an experimental test of theory using tree-hole communities. Am. Nat., 152, 510–529.

Storch, D., (2016). The theory of the nested species–area relationship: geometric foundations of biodiversity scaling. J. Veg. Sci., 27, 880–891.

Storch, D., Bohdalková, E. and Okie, J., (2018). The more-individuals hypothesis revisited: the role of community abundance in species richness regulation and the productivity-diversity relationship. Ecology Letters 21, 920–937

Thompson, G.G. and Withers, P.C., (2003). Effect of species richness and relative abundance on the shape of the species accumulation curve. Aust. Ecol., 28, 355–360.

Tuomisto, H., (2010). A diversity of beta diversities: straightening up a concept gone awry. Part 1. Defining beta diversity as a function of alpha and gamma diversity. Ecography, 33, 2–22.

Whittaker, R.H., (1960). Vegetation of the Siskiyou mountains, Oregon and California. Ecol. Monog., 30, 279–338.

Whittaker, R. J. Willis, K. J., and Field, R. (2001). Scale and species richness: towards a general, hierarchical theory of species diversity. J. of Biogeog. 28: 453–470

